# Considering founding and variable genomes is critical in studying polyploid evolution

**DOI:** 10.1101/738229

**Authors:** Xueling Ye, Haiyan Hu, Hong Zhou, Yunfeng Jiang, Shang Gao, Zhongwei Yuan, Jiri Stiller, Chengwei Li, Guoyue Chen, Yaxi Liu, Yuming Wei, You-Liang Zheng, Chunji Liu

**Author notes:** These authors contributed equally to this publication (Xueling Ye, Haiyan Hu, Hong Zhou).

## Abstract

A wide range of differences between the subgenomes, termed as subgenome asymmetry or SA, has been reported in various polyploids and different species seem to have different responses to polyploidization. We compared subgenome differences in gene ratio and relative diversity between artificial and natural genotypes of several allopolyploid species. Surprisingly, consistent differences in neither gene ratio nor relative diversity between the subgenomes were detected between these two types of polyploid genotypes although they differ in times exposed to evolutional selection. As expected, the estimated ratio of retained genes between a subgenome and its diploid donor was invariably higher for the artificial allopolyploid genotypes due likely to the presence of variable genome components (VGC). Clearly, the presence of VGC means that exaggerated differences between a donor and a subgenome in a polyploid are inevitable when random genotypes were used to represent species of either a polyploid or its donors. SA was also detected in genotypes before the completion of the polyploidization events as well as in those which were not formed via polyploidization. Considering that significant changes during and following polyploidization have been detected in previous studies, our results suggest that the influence of VGC needs to be considered in evaluating SA and that diploid donors may define changes in polyploid evolution.

## Introduction

Polyploidy, forming new species by enveloping two or more genomes within a single nucleus, is a prominent speciation process. Previous studies show that polyploidization was likely involved in the evolution of all flowering plant species (Bowers et al. 2003; Paterson et al. 2006; Jiao et al. 2011; Van et al. 2017; Ding and Chen, 2018). Understanding why polyploid species have been so pervasive and successful has attracted wide attention in the scientific community worldwide. As a result, possible effects of polyploidization on various aspects of plant biology have been intensively investigated (Feldman et al. 1997; Soltis and Soltis 1999; Li et al. 2014a; Soltis and Soltis 2016; Van et al. 2017; Edger et al. 2018; Ghanbari et al. 2019).

There are two major types of polyploids: autopolyploids and allopolyploids. The former results from whole-genome duplication (WGD) and each of them bears two or more copies of the same genome. The majority of the latter results from inter-specific or inter-generic hybridization followed by chromosome doubling, and each of them bears two or more different genomes enveloped within a single nucleus. Many important crop species, including wheat, cotton, *Brassica*, potato and banana, are typical allopolyploids.

Results from previous studies show that different species react differently to polyploidization. Wheat (Feldman et al. 1997; Uauy et al. 2017) and *Brassica* (Gaeta et al. 2007; Liu et al. 2018) are among species for which rapid and massive changes in DNA sequences were detected in newly synthesized allopolyploid genotypes. However, such changes were not detected in either cotton (Liu et al. 2001; Page et al. 2016; Wang et al. 2018) or *Arabidopsis* (Wang et al. 2006; Del Pozo et al. 2015). It is also widely believed that different subgenomes of a polyploid species have different responses to internal interactions as well as external stimuli. Such differences, termed as contrasted plasticity, subgenome dominance, or asymmetric evolution (AE), result in subgenome asymmetry (SA) or biased features in gene number, diversity or expression between the constituting subgenomes (Feldman et al. 1997; Flagel and Wendel 2010; Liu et al. 2014; Zhang et al. 2015; Wang et al. 2017; Xu et al. 2018). Possible evolutionary advantages of SA have been proposed in various species (Feldman et al. 1997; Flagel and Wendel 2010; Grover et al. 2012; Pont et al. 2013; Liu et al. 2014; Zhang et al. 2015; Kasianov et al. 2017; Wang et al. 2017; Bao et al. 2018; Xu et al. 2018).

Recent studies reveal that an individual genotype does not possess all genes existing in a species. Genes not shared by different members of a species, termed as the dispensable or variable genome components (VGC), account for about 36% in bread wheat (Liu et al. 2016; Montenegro et al. 2017), 38% in barley (Ma et al. 2019), and as high as 50% in maize (Jin et al. 2016). However, possible influence of VGC in studying polyploid evolution has not been considered. Most of previous studies reporting the existence of SA were based on the analyses of polyploid genotypes only (e.g., Pont et al. 2013; Zhang et al. 2015; Wang et al. 2017). Limited comparisons between subgenomes and their progenitors have been carried out in some studies but used only random genotypes to represent the donor species (e.g., Flagel and Wendel 2010; Li et al. 2014a). Clearly, due to the existence of VGC, using random genotypes to represent either a polyploid or its progenitor species would likely lead to over-estimation of differences between a subgenome and its donor.

Artificial allopolyploid genotypes have been obtained in different studies for several polyploid species. Different from those natural genotypes, the artificial ones all have very short time-frames of existence. In other words, the artificial genotypes have not been exposed to evolutionary selection for long thus parts of the differences between them and those natural genotypes should belong to changes accumulated during evolution in the latter. Further, the exact parents for some of the artificial allopolyploid genotypes are known thus using them to estimate shared features between a given subgenome in a polyploid genotype and its diploid donor could avoid the interference of VGC. We thus decided to use the artificial polyploid genotypes as surrogates to represent the beginning of polyploid evolution in the study reported here. Our results do not only show that considering VGC is important in evaluating SA but also suggest that changes accumulated in polyploid evolution must be defined by their founding genomes.

## Material and Methods

### Genome and RNA sequences used

Sequence data from *Brassica*, wheat, cotton and *Arabidopsis* were used in this study (Supplementary table 1). All the sequences were downloaded from the National Centre for Biotechnology Information (NCBI) Short Sequence Read Archive (SRA) database (http://www.ncbi.nlm.nih.gov/sra).

For *Brassica*, a total of 26 sets of transcriptomic data from 25 genotypes and 15 sets of genomic data from 15 genotypes were used (Supplementary table 1). The transcriptomic sequences were obtained from natural genotypes representing three allotetraploid species (*B. juncea* with a genome of AABB, *B. napus* with a genome of AACC and *B. carinata* with a genome of BBCC) and their diploid progenitors (*B. rapa* for A subgenome, *B. nigra* for the B subgenome, and *B. alboglabra* for the C genome). Sequences from artificial allotetraploids for each of the three allotetraploid species and their parental genotypes were also used. They included three artificial F1 hybrids with a genome of AC (di-haploids) and an allotetraploid derived from reciprocal hybridization between the diploid A and C donors (genomes CCAA). In addition, genomic sequences from 15 genotypes each representing the three diploid donor species were used to estimate the difference in relative genetic variation among these species. For wheat, 30 sets of transcriptomic data from 16 different genotypes were collected for this study (Supplementary table 1). They included genotypes representing natural and artificial allotetraploids (genome AABB), natural and artificial allohexaploids (genome AABBDD), their three diploid donors (*T. urartu* accession for A genome, *Ae. longissima* accession for the S genome, *Ae. tauschii* accession for the D genome), and one tatreploidploid donor (*T. durum* (genome AABB) as the maternal donor to the hexaploid wheat). The wheat genotypes also included an artificial tri-haploid (genome ABD) and a hexaploid (genome AABBDD) derived from it by chromosome doubling. It is of note that the S genome from *Ae. Longissimi* was used as the surrogate for the B subgenome donor in the artificial polyploids. Results from a recent study indicates that the B subgenome donor was likely the product from introgressions of several *Aegilops* species that yet to be identified if not already extinct (El Baidouri et al. 2017). For cotton, genomic sequences from seven genotypes were used (Supplementary table 1).

They included four natural allotetraploids, two belonging to the subspecies *G. hirsutum* and the other two belonging to *G. barbadebse.* They all share the same genome structure of AADD (Song et al. 2017). They also include one artificial tetraploid (genome AADD) and its two diploid parents (an accession of *G. arboretum* as the A genome donor and an accession of *G. raimondii* as the D genome donor).

Eighteen sets of transcriptomic data from *Arabidopsis* were also used in this study (Supplementary table 1). They were derived from one artificial and nine natural allotetraploid genotypes (all with a genome of AATT) with their two diploid subgenome donors (an accession from *A. arenosa* as the A genome donor and an accession from *A. thaliana* as the T genome donor).

### Transcriptomic sequence analyses

Based on the initial assessments using the software FastQC (version 0.11.5) (Andrews 2016), transcriptomic sequences were trimmed and filtered using the software SolexaQA++ (Cox et al. 2010). Low quality reads were removed with the criteria of Q<30 and length<50bp. As both the quality and quantity of sequences would likely influence the estimation of the numbers of genes in a genome and the percentage of shares genes between genotypes, we decided to control the quantity of sequences for each of genotype based on its relative genome size. For example, as the B subgenome or its diploid progenitor is bigger than the diploid A genome donor in wheat (Ling et al. 2013; IWGSC 2014; Avni et al. 2017), a larger quantity of the trimmed sequences was used for the former. Further, similar quantities of sequences were used for the artificial and natural genotypes (Supplementary table 1).

Following the quality and quantity controls, the selected transcriptomic sequences were assembled using Trinity (version 2.5.1) (Grabherr et al. 2011) with K-mers=25 and *de novo* assembled with a minimum length of 200bp. Redundant sequences were then removed using the cd-hit package (version 4.6.4) (Li and Godzik 2006) with the sequence similarity threshold of 95%. Coding sequences (CDS) for the assembly sequences were then predicted using TransDecoder (version 5.0.2) (https://github.com/TransDecoder/TransDecoder/releases).

In estimating genes shared between the subgenomes with their respective diploid donors, BLAST+ (version 2.7.1) (Camacho et al. 2009) was used to identify syntenic genes with a minimum E-value of 1e-5, a minimum coverage size of 100bp and a minimum identity threshold of 95%. The diploid sequences were first used as a query to BLAST against the polyploid sequences and then a reciprocal analysis was carried out by using the polyploid sequences as a query to BLAST against the diploid sequences. A custom script (Powell et al. 2017) was used to retrieve all those sequences where the polyploids or diploids gave the best hits. These sequences were used to calculate the percentages of genes shared between a given subgenome and its diploid donor.

### Genomic sequence analyses

Quality control of the genomic sequences was also carried out using the software FastQC (version 0.11.5) (Andrews 2016) and low-quality sequences were removed using SolexaQA++ (Cox et al. 2010) with Q<30 and length<50bp. The selected genomic reads were assembled using the software SOAPdenovo2-r240 (Luo et al. 2012) with the default settings based on k-mer size 63 for cotton, and k-mer size 55 for *Arabidopsis*. The GapCloser (including in SOAPdenovo2 package) was used to reduce gaps among the assembled scaffolds and the cd-hit package (version 4.6.4) (Li and Godzik 2006) was used to remove redundant sequences. For the genomic sequences from cotton, gene prediction was conducted by comparing sequences with known protein structures with DIAMOND blastx (Buchfink et al. 2015) (*E* value < 10^−10^ and bitscore > 60) based on the SWISS-PROT database (Bairoch and Apweiler 1999). The gene prediction of *Arabidopsis* was conducted by augustus (version 3.2.3) (Stanke and Morgenstern 2005) with the default settings. The genes predicted from *A. thaliana* and *A. arenosa* were used to identify shared genes in the subgenomes of the artificial allotetraploids, because sequences from the parents of the artificial allotetraploid were not available. The same methods described above for the transcriptomic sequences were used to identify genes in genome or subgenome and to calculate the percentage of genes shared between them.

For estimating the relative diversity among the three diploid donor species in *Brassica*, genome sequences were used to identify SNPs in each of them. Sequence reads were mapped to the reference genome of *B. rapa* (assembly Brapa_1.0), *B. nigra* (assembly ASM168289V1) and *B. oleracea* (assembly BOL) with BWA (version 0.7.12) (Li and Durbin 2009), after viewed and sorted by SAMtools (version 1.5) (Li 2011), duplicates were removed by samtools rmdup and SNPs were identified using SAMtools mpileup by skipping alignments with mapQ smaller than 20, finally filtered SNPs with DP < 4 and SNPs within 3 base pairs of an indel by SAMtools/BCFtools (version 1.5) (Li 2011). Genome diversity for each of the species was calculated by dividing the total numbers of SNPs with the genome size.

## Results

### Gene ratios between the subgenomes for a given species are highly consistent despite of the wide variation in predicted CDS or gene numbers among different data sets used

As expected, the numbers of predicted CDS or genes do not only vary among genotypes but also for a given subgenome among genotypes. The largest variations were found for the three polyploid species of *Brassica*, where more than two-fold differences in the number of predicted genes were detected. These differences apparently reflect differences in the quantity and quality of sequences from various sources (Supplementary table 2). However, the differences did not seem affected the estimated gene ratios between subgenomes much. In fact, the estimated gene ratios between subgenomes among various genotypes for a given species are highly consistent. It was the same subgenome which gave a lower or higher ratio of genes among different genotypes for a given species, irrespective of either transcriptomic or genomic sequences were used for the analyses. Specifically, the B subgenome is the one giving a higher ratio of genes in *B. juncea* (genome AABB); C subgenome for *B. carinata* (genome BBCC); C subgenome for *B. napus* (genome AACC); B subgenome for tetraploid (genome AABB) wheat and D subgenome for the tetraploid cotton (genome AADD) (table 1). The highly consistent gene ratios obtained between subgenomes among the different allopolyploid genotypes indicate that the use of large quantity of high-quality sequences is not critical in estimating gene ratios among subgenomes.

**Table 1.**
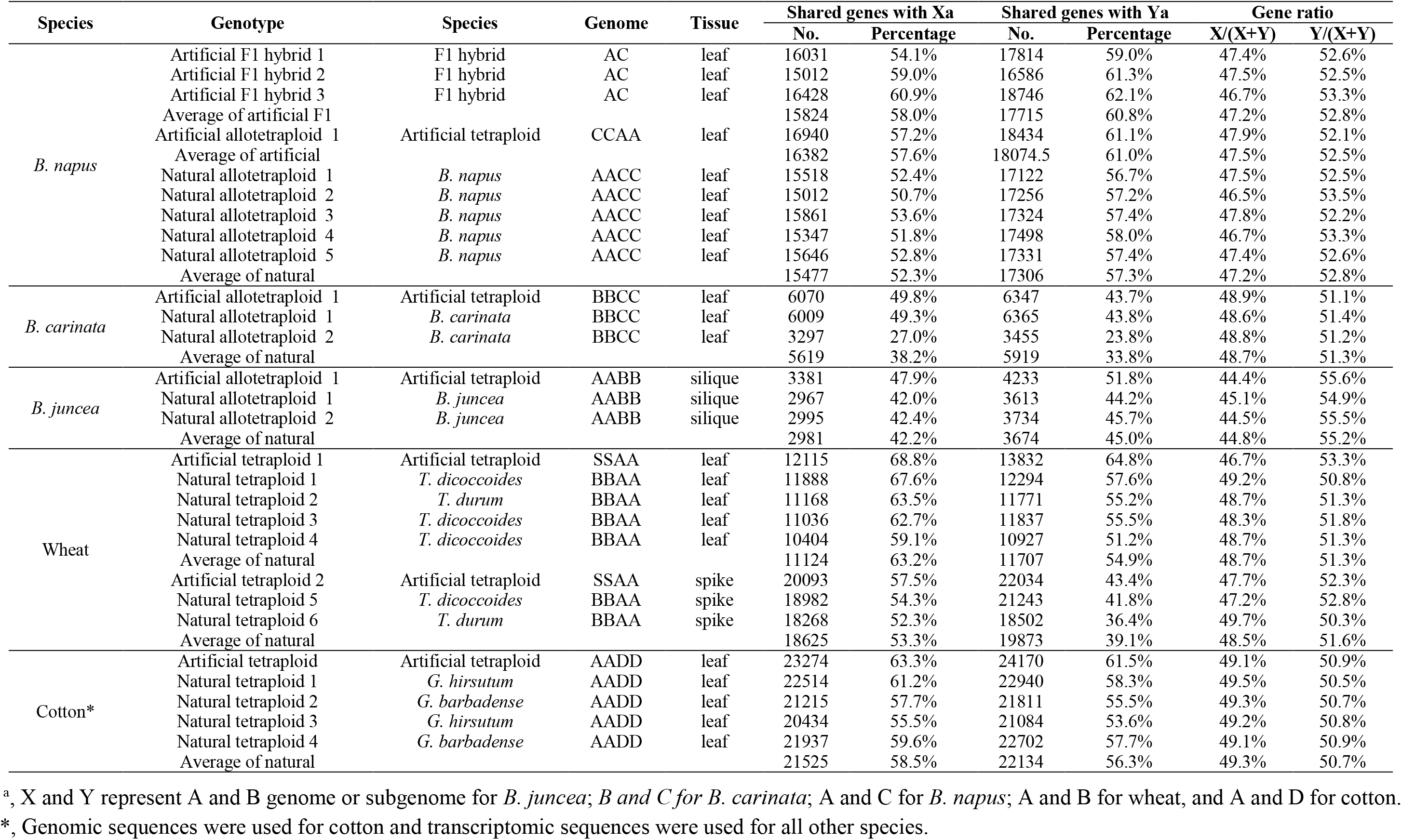
Shared genes between subgenomes and their respective progenitors and gene ratios between the subgenomes of different genotypes in different species

### Significant and consistent difference in gene ratio among subgenomes was not detected between natural and artificial allopolyploids

Compared with natural ones, the artificial allopolyploid genotypes all have very short times frames of existence. We thus decided to use the artificial genotypes as surrogates to represent the beginning of polyploid evolution, and estimated changes accumulated during evolution. Surprisingly, little difference in gene ratio was detected between the artificial and natural polyploid genotypes for any of the species studied when sequences from the same tissue was used. These included the estimations based on the use of the transcriptomic data for the four artificial and five natural allotetraploid genotypes of *B. napus* (table 1 and fig.1); for the one artificial and two natural genotypes of *B. carinata* (table 1 and fig.1), for the one artificial and two natural polyploid genotypes of *B. juncea* (table 1 and fig.1), for the six natural and two artificial tetraploid genotypes of wheat (table 1 and fig. 1). Similar results were also obtained based on the analysis of genomic sequences on cotton, although the natural genotypes of this species have a much longer history of evolution compared with either the polyploid *Brassica* or wheat (Paterson et al. 2012) (table 1 and fig. 1).

**Figure 1.**
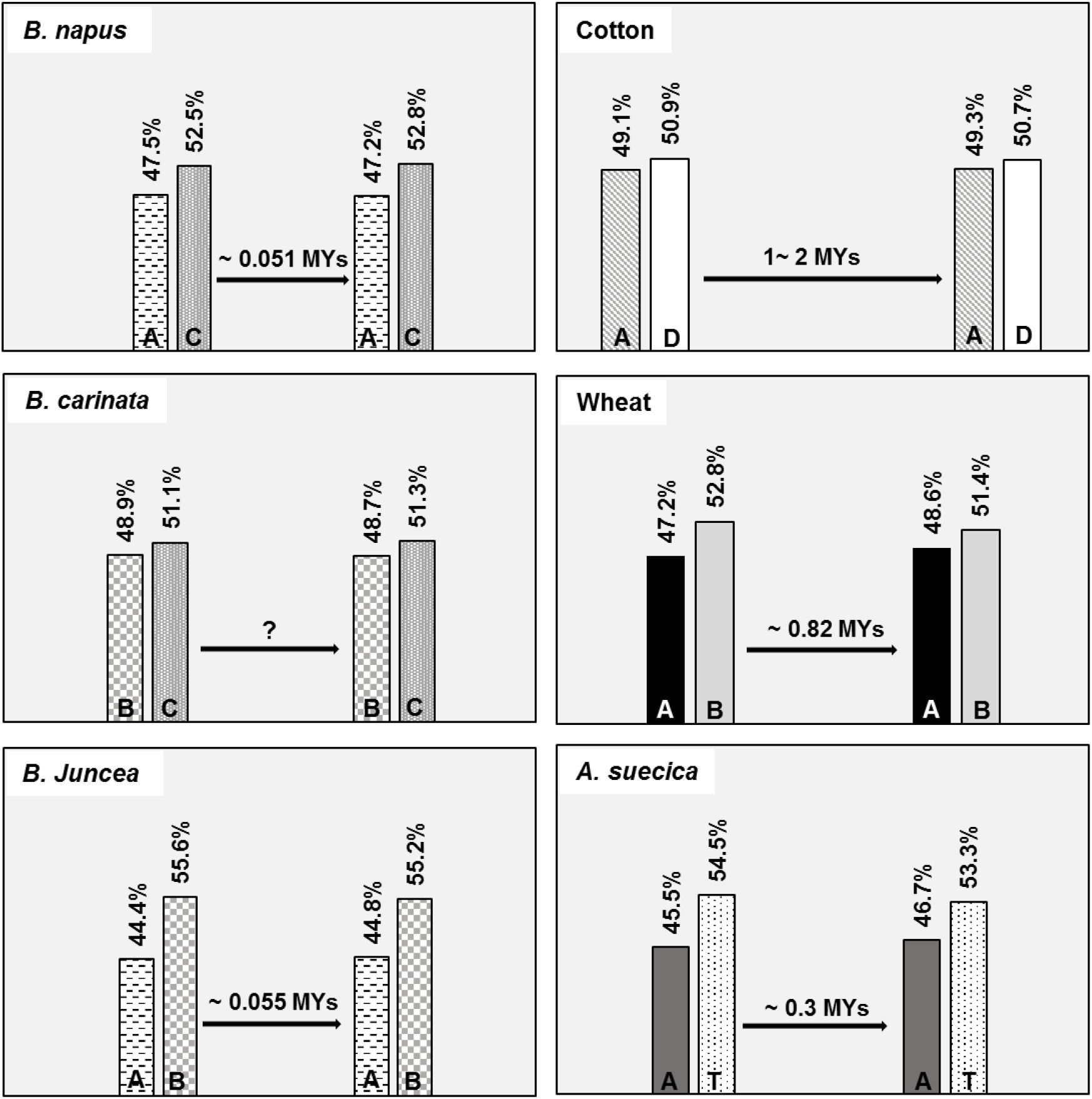
Gene ratios between the subgenomes of various polyploid species. Although there is a time difference between their formations, consistent difference in gene ratio between the subgenomes do not exist between artificial and natural genotypes for any of the polyploid species. ‘?’ indicates that the time is unknown for the species in concern, and MYs stands for million years.

### Difference in gene ratios between subgenomes was detected before polyploidization events

As mentioned earlier, the process of forming the majority of allopolyploids can be divided into two steps: inter-specific or inter-generic hybridization and chromosome doubling. Assuming that the two diploid species (one with a genome of XX and the other YY) are involved in forming an allotetraploid genotype, the first step results in the formation of a di-haploid with a genome XY, and the allotetraploid with a genome XXYY will then be formed with the completion of the second step. To examine possible changes in gene ratios among the subgenomes before and after the polyploidization event, we analysed an artificial F1 tri-haploid hybrid (genome ABD) and its corresponding allohexapolyploid genotype (genome AABBDD) based on the available transcriptome sequences from these materials. This analysis showed that the gene ratios among the subgenomes between these two genotypes were very similar (table 2). As rapid and intensive changes following polyploidisation have been reported in previous studies (Soltis and Soltis 1999; Adams et al. 2005; Soltis et al. 2015; Soltis and Soltis 2016), the results from this analysis can only be explained by the likelihood that the founding genomes define changes caused by polyploidization in the subgenomes of a polyploid.

**Table 2.** Lack of differences in gene ratio between the subgenomes of an artificial F1 hybrid and the hexaploid wheat genotype derived from it*

| Genotype (genome) | Tissue | AABB |  | DD |  | AB(D+AB) | D/(D+AB) |
| --- | --- | --- | --- | --- | --- | --- | --- |
|  |  | No. CDS | percentage | No. CDS | percentage |  |  |
| Artificial F1 hybrid (BAD ) | leaf | 25496 | 48.4% | 18467 | 49.5% | 58.0% | 42.0% |
| Artificial hexaploid (BBAADD ) | leaf | 25494 | 48.4% | 18389 | 49.3% | 58.1% | 41.9% |

\*The triploid F1 hybrid (with a triploid genome of ABD) was obtained by crossing an allotetraploid genotype (producing AB gametes) with a diploid genotype (producing D gametes). The artificial hexaploid (genome AABBDD) was derived from doubling the chromosomes of the triploid F1 hybrid.

### Gene ratios in allopolyploid genotypes not formed via polyploidization events

Different from those for the majority of allotetraploid species, the allotetraploid species of *Arabidopsis suecica* can be synthesized without the step of polyploidization. As the A subgenome donor is already a tetraploid (with a genome of AAAA), the first step in generating artificial *A. suecica* genotypes is to generate an autotetraploid genotype of *A. thaliana* by doubling its chromosomes. Then an allotetraploid with a genome of AATT can be generated by crossing the two autotetraploid genotypes. Thus, comparing gene ratios between the artificial allotetraploid genotypes with those natural ones of *A. suecica* provides another mean to test the possible effect of polyploidization on SA. Again, little difference in gene ratio between the subgenomes was detected between these two types of allotetraploid genotypes (table 3 and fig. 1).

**Table 3.** Shared genes between subgenomes and their respective progenitors and gene ratios between the subgenomes in natural and artificial *A.suecica* based on transcriptomic sequences

| Species | Genotype | Genome | Tissue | Shared gene with AA |  | Shared gene with TT |  | Gene ratios |  |
| --- | --- | --- | --- | --- | --- | --- | --- | --- | --- |
|  |  |  |  | No. | Percentage | No. | Percentage | AA/(AA+TT) | TT/(AA+TT) |
| *A. suecica* | Artificial allotetraploid 1 | AATT | leaf | 12713 | 58.1% | 15227 | 65.8% | 45.5% | 54.5% |
| *A. suecica* | Natural allotetraploid 1 | TTAA | leaf | 12071 | 55.1% | 13464 | 58.2% | 47.3% | 52.7% |
| *A. suecica* | Natural allotetraploid 2 | TTAA | leaf | 11241 | 51.3% | 13375 | 58.2% | 45.7% | 54.3% |
| *A. suecica* | Natural allotetraploid 3 | TTAA | leaf | 12004 | 54.8% | 13399 | 57.9% | 47.3% | 52.7% |
| *A. suecica* | Natural allotetraploid 4 | TTAA | leaf | 12056 | 55.1% | 13198 | 57.1% | 47.7% | 52.3% |
| *A. suecica* | Natural allotetraploid 5 | TTAA | leaf | 11389 | 52.0% | 13377 | 57.8% | 46.0% | 54.0% |
| *A. suecica* | Natural allotetraploid 6 | TTAA | leaf | 10238 | 46.8% | 13129 | 56.8% | 43.8% | 56.2% |
| *A. suecica* | Natural allotetraploid 7 | TTAA | leaf | 12110 | 55.3% | 13092 | 56.6% | 48.1% | 51.9% |
| *A. suecica* | Natural allotetraploid 8 | TTAA | leaf | 12052 | 55.0% | 13210 | 57.1% | 47.7% | 52.3% |
| *A. suecica* | Natural allotetraploid 9 | TTAA | leaf | 10873 | 49.7% | 12657 | 54.7% | 46.2% | 53.8% |
| **Average of natural** |  |  |  | 11559 | 52.8% | 13211 | 57.2% | 46.7% | 53.3% |

### Differences in relative gene diversity among the three subgenomes of the various allopolyploid *Brassica* species correlate with those among their diploid donors

To evaluate the possible relationship between a subgenome and its diploid donor, we estimated the gene diversity among the three diploid donors which contributed to the three subgenomes of the polyploid genotypes assessed. As expected, the numbers of predicted SNPs increased with the increase in quantity of sequences used. With a similar amount of sequences used, the relative gene diversity among the three diploid donors was in the order of *B. rapa* (AA) > *B. oleracea* (CC) > *B. nigra* (BB), which is the same as their order as subgenomes in the various allotetraploids (Supplementary table 2). The same order in relative gene diversity between the three subgenomes and the three diploid donors again cannot be a coincidence. Rather, they must indicate changes occurred in the subgenomes must be defined by their founding genomes.

### Difference in the ratio of shared genes between a subgenome and its diploid donor between artificial and natural polyploid genotypes

Considering the existence of VGC, the ratios of shared genes between a subgenome and its diploid donor should be higher for an artificial polyploid than for those natural polyploids. This is because that the exact diploid donors for the former are known thus influences from the VGC can be removed when conducting such an analysis. We conducted such a comparison for *Brassica*, wheat, cotton and *Arabidopsis*. As expected, the ratios shared genes between the artificial tetraploid with its two diploid parental genotypes are indeed significantly higher than those shared between the natural genotypes and those ‘random’ diploid genotypes from the progenitor species for each of these species assessed (tables 1, 3). These results reflect the importance of considering variations among individual genotypes within and between species in investigating the mechanism of polyploid evolution. In other words, the presence of VGC means that using random diploid genotypes would invariably lead to under-estimation of shared gene ratios between a subgenome and its diploid donor.

## Discussion

Polyploids are pervasive in flowering plants and SA has been reported in all allopolyploid species investigated to date. Differences in many aspects, including gene number and relative diversity, have been used to assess SA (Akhunov et al. 2003; Qi et al. 2004) and that and epigenetic variation seems to affect subgenome dominance (Hollister and Gaut 2009; Guo and Han 2014; Song et al. 2015; Cheng et al. 2016; Bird et al. 2018; Cheng et al. 2018; Ding and Chen 2018; Xu et al. 2018). Many previous studies on polyploid evolution analysed natural genotypes only and treated differences detected between subgenomes as originated from asymmetric evolution (Flagel et al. 2010; Schnable et al. 2011; Feldman and Levy 2012; Cuadrado et al. 2013; Guo and Han 2014; El Baidouri et al. 2017; Bao et al. 2018; Xu et al. 2018). To investigate possible changes in SA during evolution, we compared artificial and natural genotypes of several allopolyploid species. Different from those natural genotypes, artificial polyploids have hardly been exposed to evolutionary processes and their exact donors are known. With the believe that AE alters SA, we expected to find significantly enhanced levels of differences between the subgenomes in the natural genotypes. Surprisingly, little differences in either gene ratio or relative diversity among subgenomes were detected between these two types of allopolyploid genotypes. By examining di-haploid genotypes, we further showed that polyploidization does not seem to have dramatic influence on SA in allopolyploid genotypes. Rather, our results suggest that the differences observed between subgenomes of a polyploid seem to simply reflect those between the diploid donors of a polyploid. The less variable subgenome of a polyploid genotype was derived from the less variable diploid donor and they tend to share a higher ratio of genes. These results seem to contradict with those previous studies which have shown unequivocally that extensive changes do occur during and following polyploidization in at least some species (Qi et al. 2004; Cheng et al. 2016; Edger et al. 2017). The contradiction can be explained by the likelihood that founding genomes are critical in defining changes during polyploid evolution. In other words, considering that significant changes in various aspects including gene number (Akhunov et al. 2003; Qi et al. 2004), diversity (Adams et al. 2005) and epigenetic regulation (Wendel et al. 2000; Guo and Han 2014; Song et al. 2015; Cheng et al. 2016; Ding and Chen 2018; Edger et al. 2018) have been detected in polyploid evolution, our results suggest that the levels and directions of the changes during and following polyploidization events could be profoundly influenced by the founding genomes.

It is important to note that assaying whole gene contents for any of the genotypes used in this study is not our intention and that they are not required for estimating either gene ratios or relative diversity between subgenomes. Clearly, only expressed genes can be detected in transcriptomic analysis and the expression of many genes can be tissue-specific (Coram et al. 2008; Mittapalli et al. 2010; Fagerberg et al. 2014). However, with the use of a single set of parameters, comparing subgenomes with the large quantity of expressed genes should likely provide high-quality estimates for both gene ratio and relative diversity. This likelihood is supported by the fact that the obtained results in this study are highly consistent in that significant difference was not detected between artificial and natural polyploid genotypes for any of the species investigated with the use of transcriptomic sequences from different tissues. Considering the large number of combinations of genotypes by tissues used, the highly consistent results cannot be a coincidence. Similarly, the genomic sequences used in this study may not be adequate for high quality whole genome assembly for any of the genotypes from which they were generated. However, the results obtained from the genomic sequences again showed consistently that significant changes in neither gene ratio nor relative diversity among subgenomes were not detected between artificial and natural polyploid genotypes.

In addition to epigenetic regulations, previous studies in *Brassica* also showed that genomic exchanges including non-reciprocal homoeologous ones also affect SA (Chalhoub et al. 2014; Liu et al. 2014; An et al. 2019; Lu et al. 2019). However, similar to that found in any other species examined in this study, significant changes in SA was not detected between the artificial and natural genotypes and the differences observed between subgenomes seem to simply reflect those between the diploid donors. In fact, the triangle formed among the three diploids and their corresponding subgenomes in *Brassica* provides strong evidence showing the relationship between a subgenome in an allopolyploid and its diploid donors. The A subgenomes in either of the allotetraploids it was involved in (*B. juncea* with a genome of AABB and *B. napus* with a genome of AACC) was the one containing the lower gene ratios; the C subgenome is the one containing a higher gene ratios in both of the allotetraploids it was involved (*B. napus* with a genome of AACC and B*. carinata* with a genome of BBCC); and the B subgenome is the one containing a higher ratio in the combination of AB (*B. juncea*) but it is the one containing a lower ratio of genes in the combination of BC (*B. carinata*).

Different from many other species, the artificial allotetraploids of *Arabidopsis* (genome AATT) were not formed via typical polyploidization. They were generated by hybridizing two autotetraploid genotypes, one with the genome of AAAA and the other TTTT. In other word, no change in ploidy level was involved in the last step in obtaining the allotetraploid genotypes. However, similar to the allopolyploids of *A. suecica*, the gene ratio between the artificial tetraploids of *Arabidopsis* do not differ significantly from those of natural genotypes (table 3 and fig. 1). These results indicate that polyploidization was not be a prerequire for the differences between the two subgenomes. This likelihood is supported by that significant difference in gene ratio was not detected between the di-haploids and their allopolyploid derivatives.

Possible relationships between a subgenome and its donors are apparently not limited to gene diversity and gene ratios. For example, results from previous studies also showed that the significant difference in genome size between the subgenomes of polyploid wheats is also present between its progenitor species (IWGSC 2014; Šafář et al. 2010). Similarly, the huge difference in genome size between the two subgenomes (Zhang et al. 2015) also exists between the two progenitor species in cotton (Hendrix and Stewart 2005; Li et al. 2014b). These relationships provide further evidence suggesting that the differences between the subgenomes of the polyploid wheat are likely significantly influenced by its donors. As the exact parents for natural polyploids are unknown and VGC account for large proportions of various genomes (Jin et al. 2016; Liu et al. 2016; Montenegro et al. 2017; Ma et al. 2019), random genotypes from the donor species cannot be used to accurately estimate the relationship between a subgenome and its donor. Considering VGC or differences among individual genotypes in a species is critical in studying SA in polyploid evolution.

## Supplementary Material

**Supplemental Table 1.** Genomic and transcriptomic sequences used in this study

**Supplemental Table 2.** Numbers of SNPs detected in each of the three diploid donor species of *Brassica*

## Acknowledgments

We are grateful to Dr. Donald Gardiner for his constructive suggestions in revising the manuscript. This work was supported by the Commonwealth Scientific and Industrial Organization (CSIRO), Australia (Project code: R-10191-01). X.Y., H.Z., Y.J. and Z.Y. are grateful to the Sichuan Agricultural University and the China Scholarship Council for funding his visit to CSIRO. H.H. wishes to acknowledge Henan Institute of Science and Technology and CSC for supporting her visit to CSIRO.

## Author contributions

C.L., Y.-L.Z., Y.W., G.C., L.C. and Y.L. conceived the study; X.Y., H.Z., H.H., Y.J., Z.Y., S.G. and J.S. collected and analyzed data; X.Y., H.H., H.Z. and C.L. prepared the manuscript with contribution from other authors.

